# EpiTox: A Multi-Modular Framework for Population-Aware Off-Target Prediction Highlighting MAGEA3 Cross-Reactivity

**DOI:** 10.64898/2025.12.10.693357

**Authors:** Hoor Al-Hasani, Mazhar Ali, Stefan Krämer, Kirsten Heiss, Alexander Sinclair, Tobias Herz, Philipp Meyer, Pranav Hamde, Günter Roth, Joerg Birkenfeld, Oliver Selinger, Simon Schuster

## Abstract

Targeting intracellular tumor antigens presented by peptide human leukocyte antigen (pHLA) complexes offers a promising immunotherapeutic strategy for patients with limited treatment options. However, development of pHLA-targeted drugs such as T-cell receptor (TCR) mimic antibodies (TCRm) and TCR-based therapeutics remains challenging due to severe off-tumor and off-target toxicities arising from incomplete pHLA off-target profiling. The fatal neurological toxicities observed in the MAGEA3 TCR-T cell trial, potentially linked to unanticipated cross-reactivity with EPS8L2- and MAGEA12-derived peptides, underscore the urgent need for more comprehensive safety assessment. While current preclinical de-risking methods utilize sequence similarity searches, they lack the layered integration of genetic context, HLA binding profiles, and structured risk assessment needed to comprehensively evaluate peptide cross-reactivity. To address these limitations, we developed EpiTox, a computational multi-modular platform that systematically identifies and evaluates pHLA off-targets. EpiTox integrates proteome-wide sequence similarity, genetic context, HLA binding profiles, and layered risk assessment to provide a holistic evaluation of potential cross-reactive peptides. Using MAGEA3 as a model, EpiTox identified known toxic off-targets and predicted novel epitopes. Notably, an anti-MAGEA3 therapeutic TCRm antibody bound several newly predicted off-targets, including three SNP-derived peptides. By enabling comprehensive and predictive preclinical safety screening, EpiTox supports safer TCRm drug development and may help prevent future clinical failures.

## Introduction

The immune system uses specialized surveillance mechanisms to detect intracellular changes associated with infection or malignant transformation. Key to this process is the display of intracellular peptide fragments on the cell surface by HLA molecules. As proteins are continually degraded, their peptides are loaded onto HLA molecules and presented as peptide–HLA (pHLA) complexes, offering a snapshot of the cell’s internal state. When this repertoire shifts, such as during viral infection or oncogenic change, cytotoxic T lymphocytes (CTLs) recognize the altered pHLA complexes via their T-cell receptors (TCRs) and eliminate the affected cells (1). Because 73% of human proteins are intracellular (2), including many oncogenic drivers and tumor-associated antigens, the HLA immunopeptidome offers a large pool of targets for cancer immunotherapy. T-cell receptor mimic antibodies (TCRms) leverage this by using conventional antibody scaffolds to recognize linear peptide epitopes presented by HLA molecules rather than conformational surface epitopes. HLA class I molecules typically display 8–11 aminoacid peptides (usually 9-mers), with occasional longer peptides that bulge from the groove, creating unique design constraints. As with natural TCRs, TCRms can cross-react with different pHLA complexes that share similar surface features, raising important safety concerns due to potential off-target recognition of self-peptides (3–5). Systematic identification of physiologically relevant off-targets faces four challenges: vast search pool, population specific variants, fragmented experimental evidence, and multi-dimensional risk assessment poorly captured by single metrics. Current computational tools each address specific aspects of off-target prediction, but none yet offer a fully integrated framework encompassing the breadth of relevant biological variables. Sequence-oriented methods such as Expitope (6) and iCrossR (7) focus primarily on amino-acid identity and HLA-binding predictions, but do not explicitly incorporate population-level variation or broader biophysical properties. Biochemical similarity approaches, including dGraph (8) and sCRAP (9), leverage physicochemical descriptors, yet presently lack systematic integration of complementary evidence sources. CrossDome (10) advances the field through the incorporation of immunopeptidomics datasets, although its search space is confined to previously validated peptides, which may limit the discovery of novel off-target interactions. ARDiTox (11) incorporates SNP-derived variants, and while the conceptual strategy is outlined, detailed methodological validation and an assessment of its influence on predictive performance have not yet been presented. To address these computational and biological challenges, we developed EpiTox (Epitope Toxicity), a multi-modular platform that integrates sequence-based, structure-informed, and evidence-based methods to systematically identify and prioritize pHLA off-targets. EpiTox addresses this through 3 key innovations: [1] systematic population-specific variant integration, [2] a Bayesian framework for integrating heterogeneous experimental evidence, and [3] multi-modular ranking that captures complementary risk dimensions. We applied EpiTox to MAGEA3-KVAELVHFL, a therapeutic target associated with fatal neurotoxicity in clinical trials, to demonstrate its capability to identify both potential toxic off-targets (MAGEA12-KMAELVHFL and EPS8L2-SAAELVHFL (12, 13)) and predict novel cross-reactive peptides. Using high-throughput TCRm binding assays, we experimentally validated 538 predicted peptides, representing one of the largest systematic validation efforts for off-target prediction to date. Our results demonstrate EpiTox’s ability to recover previously reported off-targets and identify 26 novel candidate off-targets for the tested antibody that have not been described in the literature.

## Methods

### 1. Data sources and acquisition

EpiTox integrates multiple public databases at different points into its different pipelines. Datasets were first filtered according to their compatibility with each other then followed by individual preprocessing. Key parameters for each dataset are provided in (supplementary table 1). Protein sequences were fully mapped across databases by aligning both canonical and isoform variants using UniProt (14, 15) and Ensembl (16) identifiers. Unmapped sequences were excluded from the down-stream analysis. Data processing and analysis employed both established bioinformatics tools, modules and packages, and custom-built modules for EpiTox (supplementary Table 2). Key/default parameters for each analytical step are provided where applicable.

**Table 1.**
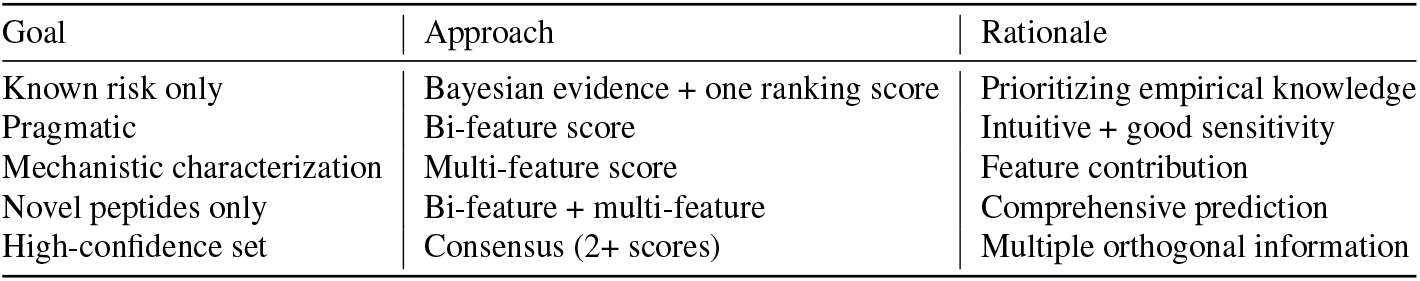
Decision matrix for experimental design.

**Table 2.** Summary of Predicted Off-Targets and MAGEA3 Known Peptides/Off-Targets.

| Predicted off-targets (n = 1454) |  |  |  |  |
| --- | --- | --- | --- | --- |
| Metric |  | High evidence<br>(n = 159, ~11%) | Moderate evidence<br>(n = 159, ~11%) | Very low<br>(n = 1195, ~82%) |
| Mean Bi-feature score (SD) |  | 0.548 (0.18) | 0.47 (0.19) | 0.34 (0.19) |
| Mean Multi-feature score (SD) |  | 0.59 (0.15) | 0.58 (0.13) | 0.57 (0.14) |
| Anchor is intact, n (%) |  | 39 (24.5%) | 26 (26.3%) | 200 (16.7%) |
| MAGEA3 known peptides/off-targets (n = 18) |  |  |  |  |
| Bi-feature | Multi-feature | N = 10<br>(validated binders, %) | N = 3<br>(validated binders, %) | N = 5<br>(validated binders, %) |
| High | High | 8 (5, 63%) | 2 (0) | 4 (3, 75%) |
| High | Moderate | 2 (1, 50%) | 1 (1, 100%) | 0 |
| Moderate | High | 0 | 0 | 1* |
| At least two scores are high |  | 10 (6, 60%) | 2 (1, 50%) | 4 (3, 75%) |

\*PP2R1B: the only known off-target in the MAGEA3 context with mismatches exclusively at anchor positions; no available experimental data was found. The low category was removed due to small size (n = 1).

### 2. Target Positional Template (TPT)

We adapted the concept of substitution matrix, that is widely used in literature to create a target-based template for target HLA-allele binding profile. For target peptide T (length *L*), all single-residue substitutions (*L* × 19) were generated, and predicted HLA binding affinities were obtained with NetMHCpan (17). For position *l*, the mean value across substitutions 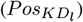 was used to derive three complementary metrics:

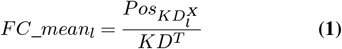

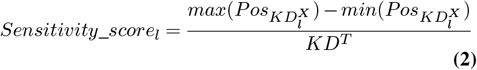

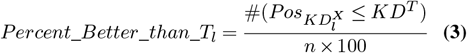

where 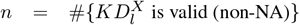 Positions in the top 30% for both *FC*_*mean* and Sensitivity were designated as anchors, those in the top 30% for *P ercent*_*Better*_*than*_*T* were considered permissive, and all others were classified as neutral. The substitution matrix provides a model-derived positional template to describe mismatch patterns between the target peptide and locally similar off-target sequences, serving as a relative NetMHCpan-based sensitivity profile rather than an experimental binding map.

#### Off-Target mismatch profile

Each off-target peptide was analyzed by projecting its mismatches onto the TPT classes: **Anchor only/mismatch**: One or more mismatches at anchor positions exclusively, potentially altering peptide-MHC complex structure (pHLA). **Backbone-only/intact**: Anchor positions conserved but exposed positions differ, maintaining presentation but potentially avoiding antibody recognition. **Both**: the mismatch is at least in one anchor and one backbone position.

### 3. EpiTox architecture

EpiTox workflow-kit consists of multiple individual workflows implemented with Snakemake (18), each designed to perform specific tasks (Fig. 1). The module’s name reflects the main task or the dominant task of the module. The different modules serve two functions, off-targets identification and peptides annotation.

**Fig. 1.**
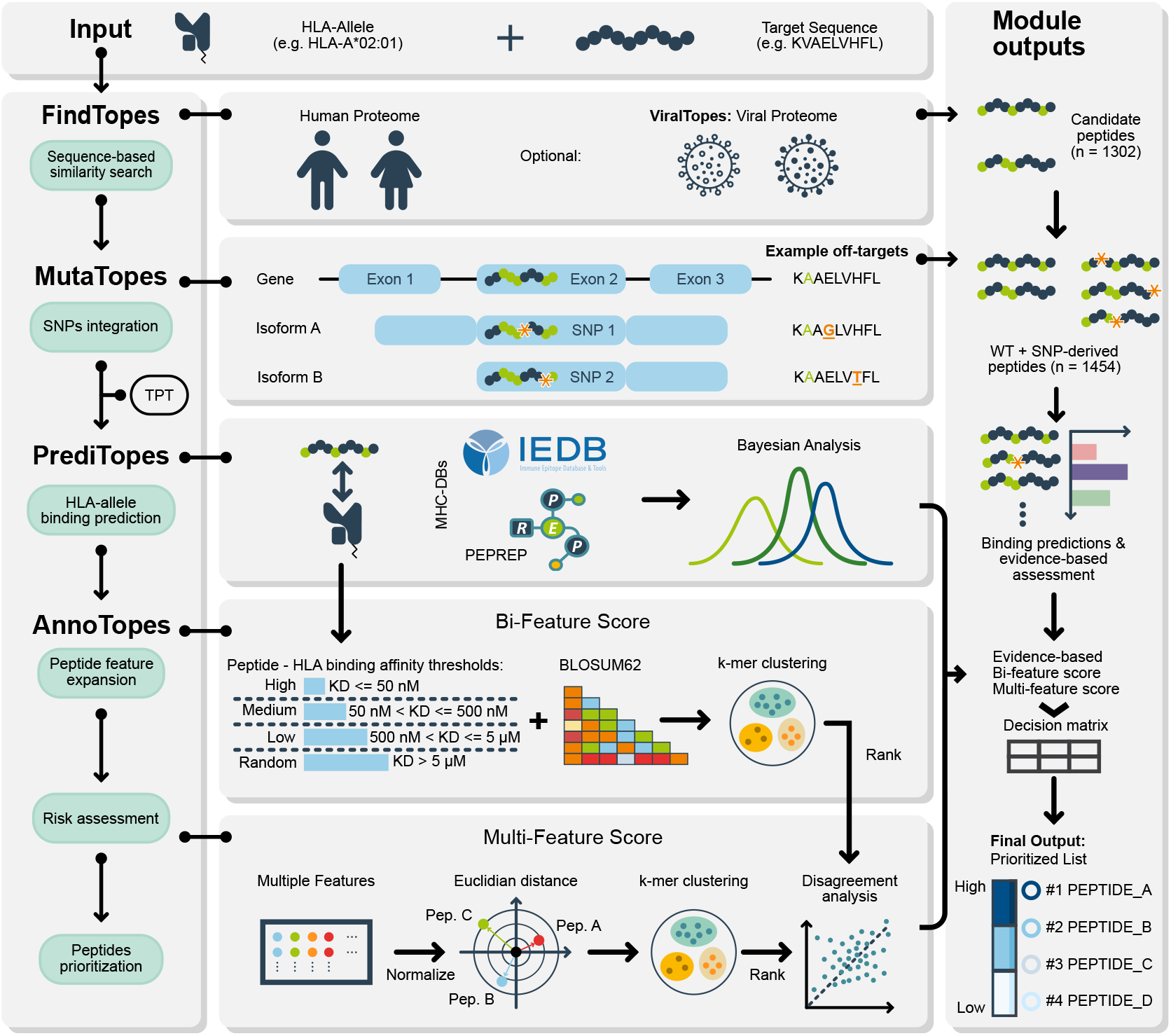
EpiTox Workflow for Off-Target Peptide Prediction: EpiTox uses five computational modules to assess off-target risks. From an input peptide sequence, FindTopes searches the human proteome for similar sequences (ViralTopes extends this to pathogen proteomes). MutaTopes generates SNP-derived peptides from population variants. PrediTopes uses NetMHCpan to predict peptide-HLA binding and performs Bayesian risk assessment using IEDB and PEPREP. AnnoTopes combines predictions with physicochemical properties to generate two ranking metrics: a Bi-feature score (sequence similarity plus HLA-binding affinity) and a Multi-feature score (eight physicochemical parameters via target-template comparison). The workflow outputs a prioritized list of potential off-target peptides for TCR-like therapeutic development.

#### 3.1. FindTopes: off-targets identification module

FindTopes identifies potential off-target candidates through complementary sequence similarity searches: BLASTp (19)(using BLO-SUM62 and PAM30 matrices) captures biologically meaningful relationships through evolutionary-informed scoring and gapped alignments, while PEPMatch (20) provides orthogonal exact k-mer matching. This dual strategy balances sensitivity and specificity by detecting both conservative substitutions that may preserve cross-reactivity and exact local sequence matches. FindTopes applies three customizable filters, Levenshtein distance, peptide length, and gap. Results shown in this study were generated with Levenshtein distance ≤ 4, length cutoff = 9, so that all off-targets are of the same length as the target, and finally only peptides without gaps were included in the downstream analysis. The generated peptides list in this module has the complete mapping of genomic and proteomic location coordinates.

#### 3.2. MutaTopes: SNPs integration

SNPs were extracted according to the chosen population. In this study SNPs were filtered by European population according to gnomAD’s annotations (FIN + NFE)^1^(21), quality and allele frequency (default ≥ 1%). Genomic coordinates for all FindTopes-identified peptides are extracted from reference annotations GRCh38. These coordinates are intersected with the selected SNPs (gnomAD v2.1.1 leftover to GRCh38) to identify variants falling within or adjacent to peptide-regions. SNPs meeting the coordinate criteria are transformed to the protein level using VEP (22), ensuring isoform-SNP mapping is maintained and retained for subsequent analysis. For each isoform-SNP, variant peptides are generated by applying amino acid substitution to all overlapping peptide windows (in this study =9). SNP-derived peptides are filtered using edit distance cutoff < 5 and duplicate removal so that only unique isoform-peptide are kept.

#### 3.3. PrediTopes: HLA binding prediction and experimental evidence

Affinity and presentation score are predicted using NetMHCpan. Peptides experimental annotations were then fetched from IEDB (23) and PEPREP (supplementary table 1, supplementary section: PEPREP), preprocessed and harmonized for the downstream analysis. Starting from a neutral prior probability (50%), we update confidence based on available evidence using Bayes’s theorem:

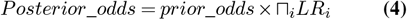

Where *Prior*_*odds* = *P* (*H*)*/P* (¬*H*) = 0.5*/*0.5 = 1, and *LR*_*i*_ represents the likelihood ratio for evidence type *i*. The posterior probability was then derived as:

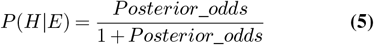

Protein-peptide pairs were classified into four evidence groups based on the probabilities: High (≥ 80%), moderate (50-79%), low (30-49%), and very low (*≤* 30%)

##### Likelihood ratios logic

Likelihood ratios (LRs) were assigned to each evidence type based on their relative strength and biological relevance, with values ranging from 0.3 (predicted-only peptides) to 12 (strong binding to target allele) (Supplementary Table 4). LR selection was guided by the following principles: [1] strong experimental evidence in the target context (same HLA allele, normal tissue, multiple studies) received high LRs (8-12), [2] moderate evidence (partial context match) received intermediate LRs (4-6), and [3] predicted-only peptides received (LR = 0.3). Sensitivity analysis confirmed that these LR values provided appropriate discrimination; the framework assigns 23% confidence to predicted-only peptides and 92% to experimentally validated peptides when starting from a neutral 50% prior (Supplementary Figure 1). This 69-percentage-point spread ensured clear separation between confidence categories while allowing realistic evidence accumulation when multiple factors were present.

#### 3.4. AnnoTopes: off-targets annotation and ranking

##### Bi-feature score

The score prioritizes off-targets based on similarity to the target, using BLOSUM62 substitution matrix (*b*), and its affinity to HLA-allele (*a*). The score is bounded [0-1], the higher the score is, the higher the risk is, i.e., highly ranked peptides are more likely to be relevant off-targets and therefore should be included in downstream analysis. The score function (*S*) is defined as:

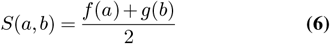

where affinity is defined as *f* (*a*) = {1, 0.66, 0.33, 0} for *a* ∈ { [0, 50), [50, 500), [500, 5000), [5000, ∞)} to maintain biological relevance and

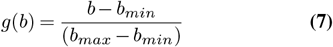

where *b*_*min*_, *b*_*max*_ are the minimum and maximum BLO-SUM62 value respectively. The final peptide classification is then estimated using supervised machine learning method, k-means with 3 clusters (EpiTox’s default clustering mode): *peptide*_*rank*_ = *kmeans*_3_(*S*) → {*Low, M oderate, High*}

##### Multi-feature score

Eight physiochemical properties for each peptide were computed: Five global features (GRAVY, isoelectric point [pI], net charge at pH 7.4, aromaticity, and aliphatic index) and three position-specific features corresponding to TCR-facing backbone residues (backbone GRAVY, backbone charge, and backbone aromaticity) at previously defined positions according to target-template (Methods 2). The choice of features was guided by observations from internal analyses on related datasets, where these properties appeared to show meaningful variability and potential relevance to pHLA and structural characteristics. All features were computed and z-score normalized. Target anchor & backbone positions were determined in the TPT (Methods 2), and the associated off-target positions classification was determined after obtaining the final list from MutaTopes. Features were weighted based on mismatch location relative to HLA anchors, these weights within each category were normalized to sum to 1. When anchors are intact, HLA presentation is maintained and TCR recognition of exposed backbone residues dominates; when anchors are disrupted, global properties affecting presentation become more relevant. Therefore, the weights are Anchor intact (backbone mismatches only): Backbone features weighted 3.0 ×, global features 1.0 ×; Anchor mismatch (backbone intact): Global features weighted 2.0 ×, backbone features 1.0 ×; Both (mismatches everywhere): Global features 1.0 ×, backbone features 1.5 ×.

Multi-biophysical distance from the target peptide was computed as weighted Euclidean distance:

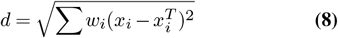

where *x*_*i*_ and 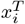 are normalized feature values for off-target and target peptides, and *w*_*i*_ is the conditional weight for feature *i*. Peptides were ranked by ascending distance (rank 1 = most similar).

#### 3.5. ViralTopes: module for pathogen derived off-target identification

ViralTopes extends FindTopes logic to pathogen genomes (e.g., HIV) using identical criteria. This optional module was not applied in the MAGEA3 case study.

### 4. Validation Candidate Selection

To accommodate the inherent complexity in predicting potential off-target peptides, we provide multiple scoring strategies rather than relying on a single metric. The choice of approach should be guided by the experimental goal, as summarized in Table 1.

### 5. Peptide–HLA array production and SCORE measurements

Peptide–HLA array production and SCORE measurements were performed as described previously (24). Peptides were purchased from Peptides & Elephants GmbH (Hennigsdorf, Germany), and biotinylated stabilized HLA-A*02:01 variant was used (25). The antibody was measured in its Fab format to ensure a 1:1 interaction. The binding curves were fitted and analyzed using an internal version of anabel and a 1:1 Langmuir binding model (26).

## Results

We developed EpiTox to enable holistic and systematic framework for assessing pHLA off-target risk assessment in drug discovery. Off-target identification is inherently multifaceted: It requires surveying the antigen landscape, evaluating cross-reactivity potential, and interpreting biological relevance. No single computational strategy can capture this complexity. To address this, EpiTox decomposes the problem into five complementary modules: FindTopes, ViralTopes, MutaTopes, PrediTopes, and AnnoTopes, each tackling a specific computational challenge (Fig. 1, Methods 3). Together, these modules form an integrated workflow that progresses from broad off-target discovery to biological prioritization. To illustrate EpiTox in practice, we used the MAGEA3 target peptide (KVAELVHFL) as a representative case study. This example demonstrates how the tool’s modular architecture enables systematic identification, classification, and prioritization of potential off-targets for an anti-MAGEA3 TCRm. These results highlight the peptide’s cross-reactivity potential but do not imply general performance across all targets.

### FindTopes and MutaTopes: Sequence-based off-Target prediction

We first derived a MAGEA3 target positional template using the scores introduced in TPT. The analysis confirmed positions P2 and P9 as classical anchor positions, and P1 as a secondary anchor position (Supplementary Table 5, Supplementary Figure 2), consistent with the established HLA-A*02:01 binding motif. In contrast, positions P3 & P5 were classified as permissive, tolerating 60-70% of amino acids substitution while maintaining the binding affinity within 2-fold of wildtype. FindTopes and MutaTopes identified 1454 putative sequence similar off-targets for MAGEA3-KVAELVHFL, of which 1304 were wildtype sequences and 150 were SNP-derived peptides (Fig. 2.A). SNP-derived peptides exhibited varying sequence divergence: one peptide contained two mismatches, seven contained three mismatches, and 142 contained four mismatches. Five SNP-derived peptides contained irregular symbols resulting from stop codon annotations at the protein level. Of the 19 off-targets previously disclosed in patents and studies, FindTopes successfully identified 18 (Fig. 2.B, Supplementary Table 3). The one missing off-target, MAGEA5-KVADLIHFL, was excluded due to quality controls applied during data aggregation (Methods 1). According to the TPT analysis (Supplementary Table 5), around 19% of off-targets have sequence mismatches within the backbone positions only (Fig. 2.C). Almost 81% of the predicted off-targets have their sequence variation both in anchor and permissive positions. Only three peptides have mismatches exactly at the anchor positions, MAGEA12-KMAELVHFL with one mismatch at P2, EPS8L2-SAAELVHFL with two mismatches at P1 & P2 and PPP2R1B-GIAELVHFS with three mismatches at all classified positions P1, P2 & P9. SNP-derived peptides follow the overall mismatch classification pattern, with sequence alterations in both peptide backbone and anchor positions (n = 125, 83%) and a smaller proportion showing intact anchor positions (n = 25, 17%). Finally, several of the top peptides with the lowest Levenshtein distance were, as expected, from the MAGE family (Supplementary Table 6) including MAGEA9, a perfect match to the target sequence (Levenshtein distance = 0).

**Table 3.** Overview of features supported by EpiTox and other state of the art tools.

| Metric/Outcome | EpiTox | Typical tools* | Note |
| --- | --- | --- | --- |
| **Discovery and Validation** |  |  |  |
| Peptides experimentally tested | 538 (plus target) | 20-50 | 10 – 25× scale |
| Known off-targets recovered | 18/19 | Not benchmarked | 95% recall |
| Hit rate | 9.7% | unknown | Enrichment over proteome baseline |
| Novel cross-binders | 26 | Rarely reported | Discovery focus |
| SNP-derived binders validated | 3 | Not addressed | Population risks |
| **Candidate pool** |  |  |  |
| Off-target candidates identified (MAGEA3) | 1454 | Varies | Comprehensive |
| Wild-type peptides | 1304 | Typical | Standard |
| SNP-derived peptides (population specific) | 150 | 0 | Unique |
| **Evidence integration** |  |  |  |
| Database source integrated | IEDB, PEPREP | IEDB or MS data** | Comprehensive search |
| MS immunopeptidome coverage | IEDB and PEPREP | Direct MS pool*** | Some tools are MS focused |
| Protein-specific evidence tracking | Yes | No | Context-aware |
| Graduated confidence categories | 4 tiers | Binary or score | Stratification |
| **Multi-dimensional assessment** |  |  |  |
| Complementary scoring methods | 3 | 1-2 | Comprehensive |
| Binder coverage | 36/36 | Partial to the experimental selection | 100% coverage |
| Single method coverage range | 51-95% | Not reported | Complementary |

\*representative comparison to Expitope, CrossDome, ARDiTox, iCross based on published literature.
\*\* CrossDome and similar tools leverage large MS immunopeptidome datasets from HLA elution studies.
\*\*\* some tools (e.g., scrap) built directly on massive MS immunopeptidome own pool.

**Fig. 2.**
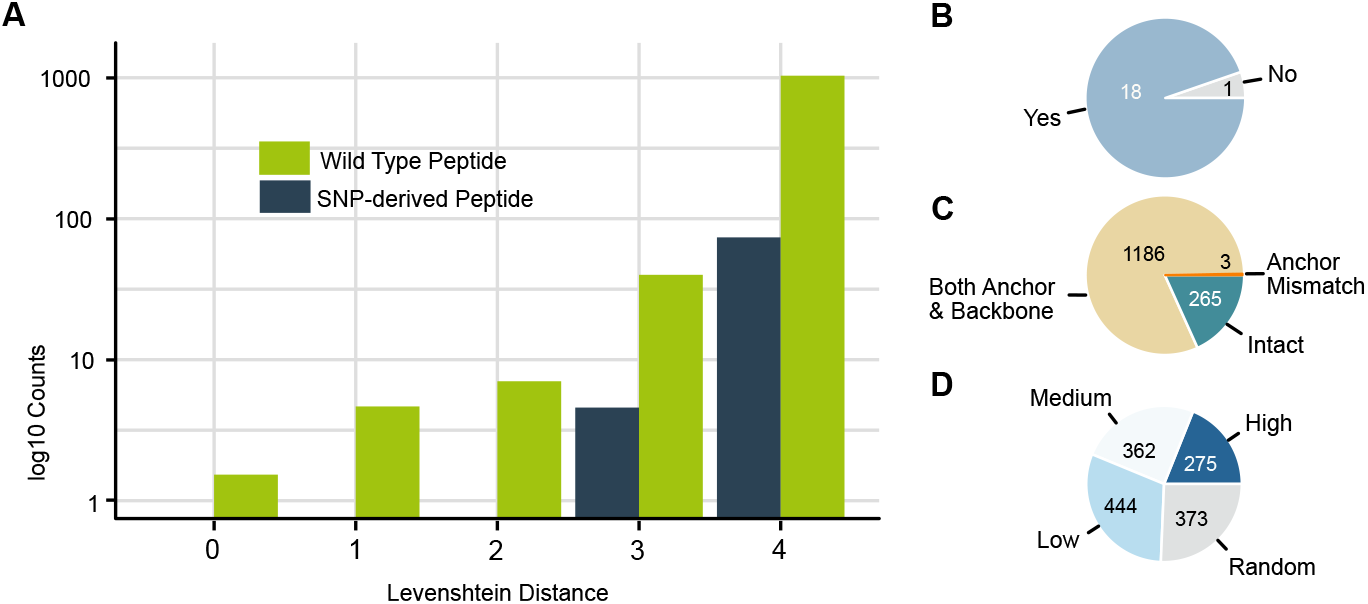
FindTopes and MutaTopes enable comprehensive identification of reference and SNP-derived off-target peptides: Selected output from the EpiTox-FindTopes and -MutaTopes modules using MAGEA3-KVAELVHFL as input. (A) Bar chart of peptides identified by FindTopes and MutaTopes in the human proteome, distributed according to their edit distance from the target. Two categories are shown: wild-type peptides and SNP-derived peptides. (B) Number of literature-known peptides from previous patents and studies identified by EpiTox-FindTopes; only one peptide was missed due to quality checks during data aggregation. (C) Position mismatch classification of predicted peptides according to the target binding profile. (D) Peptides classified by PrediTopes according to predicted HLA-A*02:01 binding affinity (NetMHCpan); approximately 50% of peptides exhibit moderate to high binding affinity.

### PrediTopes: Peptide binding prediction and Evidence-based assessment

Once off-target peptides are identified by the previous two modules, PrediTopes generates two types of characterization: [1] binding affinity predictions that serve as input feature for composite scoring in AnnoTopes, and [2] contextual evidence derived from a Bayesian model harnessing MHC database(s), which is used directly in downstream analysis. The context-aware Bayesian framework integrates multiple lines of evidence to generate confidence scores for peptide-HLA pairs. The framework addresses two key challenges: [1] maintaining protein-specific evidence tracking to prevent inappropriate validation transfer across homologs, and [2] integration of experimental validation status with computational predictions in a probabilistically sound manner (Methods 3.3).

### Binding affinity prediction for MAGEA3 off-targets and HLA-A*02:01

For each peptide identified by the two previous modules (FindTopes and MutaTopes), HLA-allele binding affinities and presentation were predicted (Fig. 2.D). Accordingly, 277 peptides were strong binders (mean KD = 28.3 nM), 444 moderate binders (mean KD = 179 nM), low binders (mean KD = 1999 nM), and 373 random binders (mean KD = 14960 nM). The predicted affinity exposed the largest variation in peptides with mismatches at both anchor positions and backbone (Supplementary Figure 3). Moreover, the three peptides with mismatches at anchor positions showed different binding: While MAGEA12-KMAELVHFL and EPS8L2-SAAELVHFL both have high predicted affinity (KD = 10.8, 29.5 nM respectively) showing a similar level to those who have intact anchor positions; PPP2R1B-GIAELVHFS was predicted to have moderate affinity (KD = 1313.3nM).

### Evidence-based assessment of MAGEA3 predicted off-targets using PrediTopes

A total of 286 peptides, representing 259 distinct peptide–protein pairs, were found in the MHC databases (IEDB and PEPREP) (Fig. 3A), upon inspection, peptide-level coverage distributions were right-skewed across all dimensions (Fig. 3.B). Mean values consistently exceeded medians: observations (8.49 vs. 3.0, 182.9%), diseases (2.57 vs. 1.0, 156.8%), and studies (4.22 vs. 2.0, 111.2%). This pattern indicates that while a minority of peptides are extensively characterized, the typical peptide exhibits limited coverage, appearing in a single disease, tested with two HLA alleles, and examined in two studies. PrediTopes assigned evidence-based risk scores through Bayesian integration of experimental MHC data (Methods 3.3). Of the 1454 candidates, 159 peptides (11%) were classified as high evidence (mean posterior probability = 99.7%, SD = 0.19), 99 (7%) peptides as moderate evidence (mean = 64.3%), 1 as low evidence (31.0%), and 1195 peptides (82%) as very low evidence (mean = 23.1%) (Fig. 3.C). The discrete tier structure reflects the categorical nature of available database evidence. Peptides with identical evidence profiles yield identical posterior probabilities (e.g., 99 peptides at 64.3%, all supported by predicted + intermediate binding and Class I data). Because the evidence is uneven across candidates, these scores are best interpreted as relative, evidence-based rankings for prioritization rather than as calibrated probabilities of immunogenicity. Notably, all SNP-derived peptides fell into the very low evidence tier, illustrating that limited experimental data translates directly into lower posterior rankings.

**Fig. 3.**
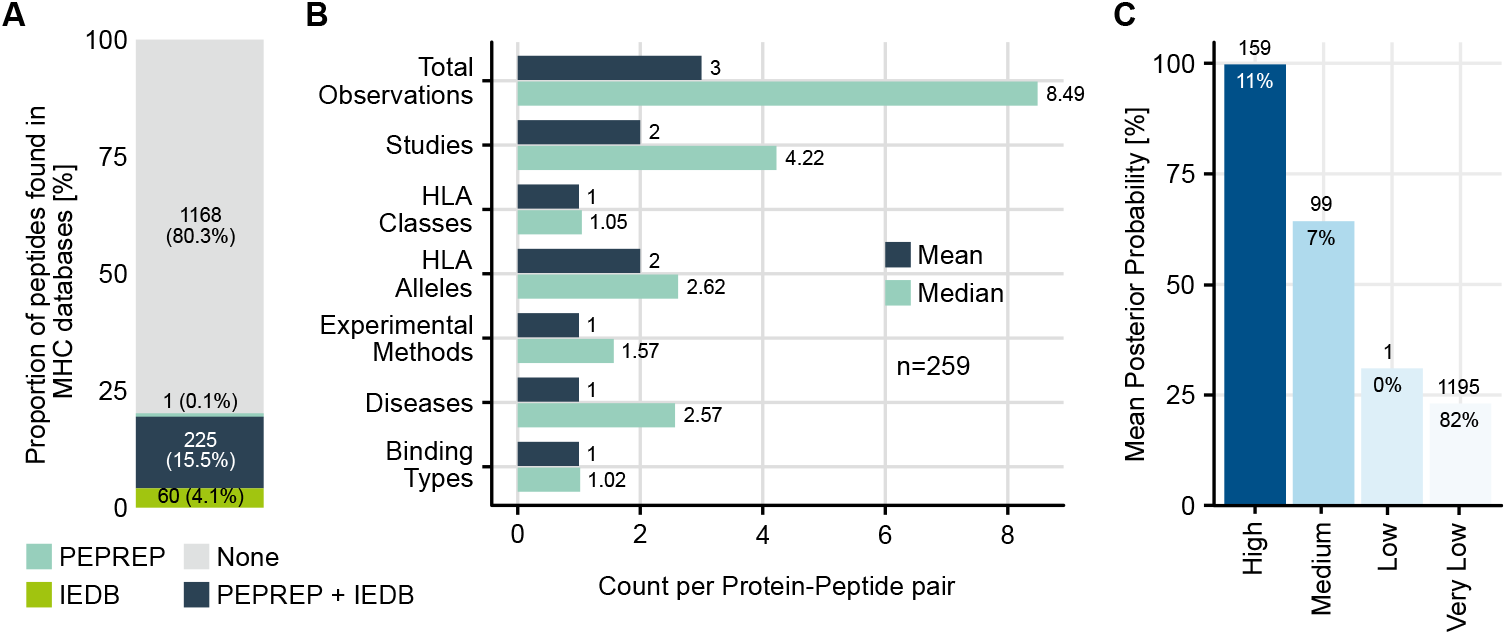
Contextual evidence of MAGEA3 off-targets derived by PrediTopes Bayesian framework: (A) peptides with available experimental data according to PEPREP (in-house database for pHLAs) and IEDB. Accordingly, 80% (1168 of 1454) of the predicted peptides have no previous record. (B) Comparison of mean and median data coverage across 259 unique protein-peptide pairs (2,198 total observations). (C) peptides classification according to experimental evidence: four evidence tiers produced by the Bayesian framework, posterior probability shows appropriate separation.

### Sequence-based evidence aggregation and protein context provided by PrediTopes are critical for accurate peptide off-target assessment

We examined 94 peptide sequences appearing in multiple proteins within our dataset to assess whether sequence-identical peptides exhibit protein-context-dependent experimental validation. Most sequence-identical peptides (84/94, 89%) lacked experimental validation for any protein source. Among the 10 peptides with available data, four (40%) exhibited differing confidence classifications depending on the associated protein (Supplementary Table 7). KVDEAVAVL received high evidence classification for PABP1/PABP4 but very low for PABP3. AVHELGHAL showed moderate evidence for MMP14 but very low for MMP15/16/24. These results demonstrate that experimental validation is protein-context specific and non-transferable across peptide homologs.

### Contextual evidence from PrediTopes distinguishes sequence-identical therapeutic target from potential off-target

MAGEA3 (P43357) and MAGEA9 (P43362) share an identical peptide sequence. We compared their evidence profiles using PrediTopes’ context-aware framework. PrediTopes assigned substantially different likelihood ratio products (10000 vs 270, representing a 37-fold difference) based on distinct experimental contexts. MAGEA3 (P43357) showed strong HLA-A*02:01 binding in normal tissue assays, while MAGEA9 (P43362) demonstrated intermediate binding with the same allele exclusively in cancer contexts. Both peptides achieved High evidence classifications, but the cumulative evidence strength differed markedly, reflecting their distinct experimental validation profiles.

### AnnoTopes: Integrated annotation and feature-based off-target ranking

Peptide cross-reactivity depends on multiple features that may not be captured by a single metric. We therefore implemented three complementary scoring strategies to provide multidimensional assessment: Contextual evidence from PrediTopes, Bi-feature scores, and Multifeature scores from AnnoTopes (Methods 3.4). These metrics capture both overall peptide composition and TCR-facing structural characteristics.

### Feature-driven Ranking of MAGEA3 predicted off-targets

#### Ranking MAGEA3 predicted off-targets with Bi-feature score

We first applied a minimalistic two-component score combining sequence similarity (BLOSUM62) and predicted MHC affinity (Methods 3.3). This approach identified three risk categories: High risk (n=299, median score = 0.64), moderate risk (n=557, median score = 0.44), and low risk (n=598, median score = 0.16) (Fig. 4.A). SNP-derived peptides were also represented across all risk categories: high risk (n=26, median score = 0.67), moderate risk (n=59, median score = 0.44), and low risk (n=65, median score = 0.14).

**Fig. 4.**
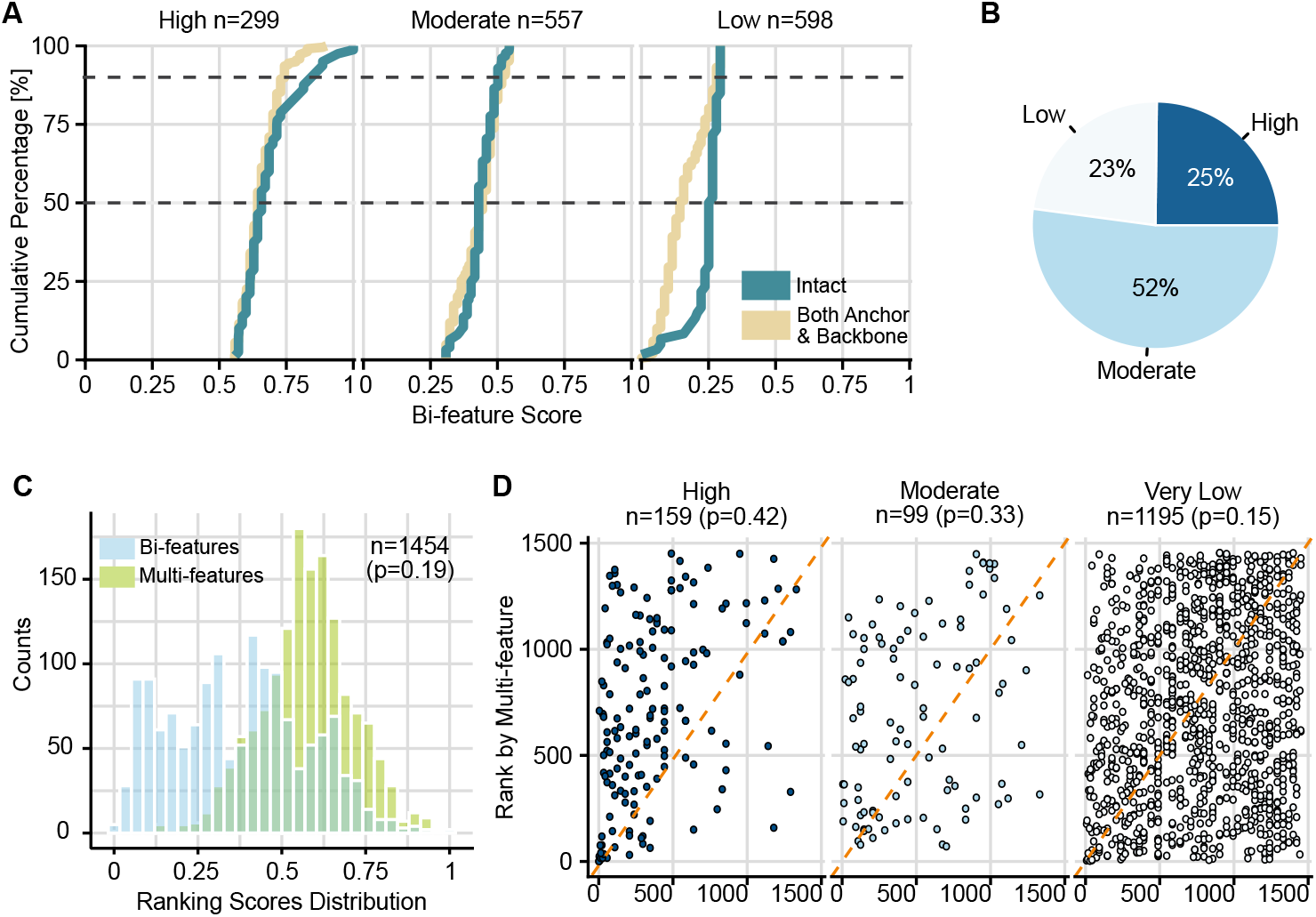
Predicted MAGEA3 off-targets ranked by AnnoTopes Bi-feature and Multi-feature scores: (A) Cumulative Distribution Function (CDF) of Bi-features score stratified by risk rank (High, Moderate, Low) according to k-mean method, and anchor status (Intact, Both Anchor, and Backbone). Peptides with mismatches at anchor positions (n=3) were excluded due to small sample size. Dashed lines indicate 50th and 90th percentiles. (B) Multi-feature score categories according to k-mean method, accordingly 25% of the peptides (n = 361) were considered highly similar to the target by sequence properties. (C) histograms of Bi-features and Multi-features scores across all off-target peptides. Low correlation (*ρ* = 0.19) demonstrates orthogonal risk dimensions. (D) Scatter plot comparing the ranked Bi-features and Multi-features scores stratified by contextual evidence-levels, peptides in category Low (n =1) were excluded due to small size. High evidence panel, known space, shows more structured distribution with reasonable agreement between Bi-and Multi-feature scores. Moderate evidence panel shows more scattered and wider spread from diagonal. very low evidence, unknown space, highly dispersed with off-diagonal scatter as the two scores show more independence.

Peptides classified as high-risk exhibited both strong predicted HLA binding affinity to HLA-A*02:01 (median affinity: 30 nM) and moderate to high BLOSUM62 sequence similarity to MAGEA3 target (median similarity score: 21; self-similarity = 44). Two-thirds (2/3) of peptides with mismatches at anchor positions demonstrated strong MHC binding affinity (KD ≤ 30 nM), comparable to intact anchor peptides. We next examined whether the Bi-feature score effectively prioritized the 18 previously reported/validated offtarget peptides (Fig. 2.B, Supplementary Table 3). Among these known peptides, 17 were classified as high risk and 1 as moderate risk. MAGEA9 received the highest rank score (1.0) due to identical sequence with the target. EPS8L2 and MAGEA12, previously implicated in fatal neurotoxic cases, received scores of 0.90 and 0.87, respectively, reflecting reduced similarity and affinity relative to MAGEA3. PPP2R1B, the only peptide with all three mismatches confined to anchor positions, was assigned a moderate risk score (0.47).

#### Ranking MAGEA3 predicted off-targets with Multi-feature score

To capture dimensions beyond sequence and binding, we computed eight physicochemical properties weighted by mismatch location (Methods 2, 3.4, Supplementary Figure 4). Feature weights were adjusted based on anchor and backbone integrity. K-means clustering (k=3) of multifeature distances yielded three categories: High similarity (n=361, mean distance = 0.75), moderate similarity (n=758, mean = 0.58), and low similarity (n=335, mean = 0.39) (Fig. 4.B). Among the 18 known off-targets, 15 were classified as high similarity and three as moderate. PPP2R1B, with three mismatches exclusively at anchor positions, was ranked as moderate risk by Bi-feature (0.47) but high similarity by Multi-feature (0.78). In contrast, two peptides with intact anchors but backbone mismatches, MAGEA6KVAKLVHFL and DDX28-KVAELVHIL, showed the opposite pattern: high risk by Bi-feature but moderate by Multifeature.

#### AnnoTopes and PrediTopes scores complement each other

We compared three scoring approaches: PrediTopes Bayesian framework for evidence-level assessment, Anno-Topes Bi-feature score for functional assessment, and Anno-Topes Multi-feature score for sequence similarity. Bi-feature and Multi-feature scores showed weak overall correlation (Spearman *ρ* = 0.19) (Fig. 4.C, Supplementary Figure 5). However, the relationship varied substantially across contextual evidence categories (Fig. 4.D). Peptides with high experimental evidence (n=159) showed moderate concordance between scoring metrics (Spearman *ρ* = 0.42) with representation across the full similarity spectrum. Peptides with moderate evidence (n=99) displayed greater scatter (*ρ* = 0.33), while peptides with very low evidence (n=1195) showed the weakest correlation (*ρ* = 0.15) and greatest scatter. High evidence peptides exhibited higher Bi-feature scores (mean 0.548, SD = 0.188) compared to very low evidence peptides (mean = 0.341, SD = 0.192) (Table 2). In contrast, Multifeature scores remained almost consistent across evidence levels (High: 0.586, moderate: 0.583, very low: 0.572).

**Fig. 5.**
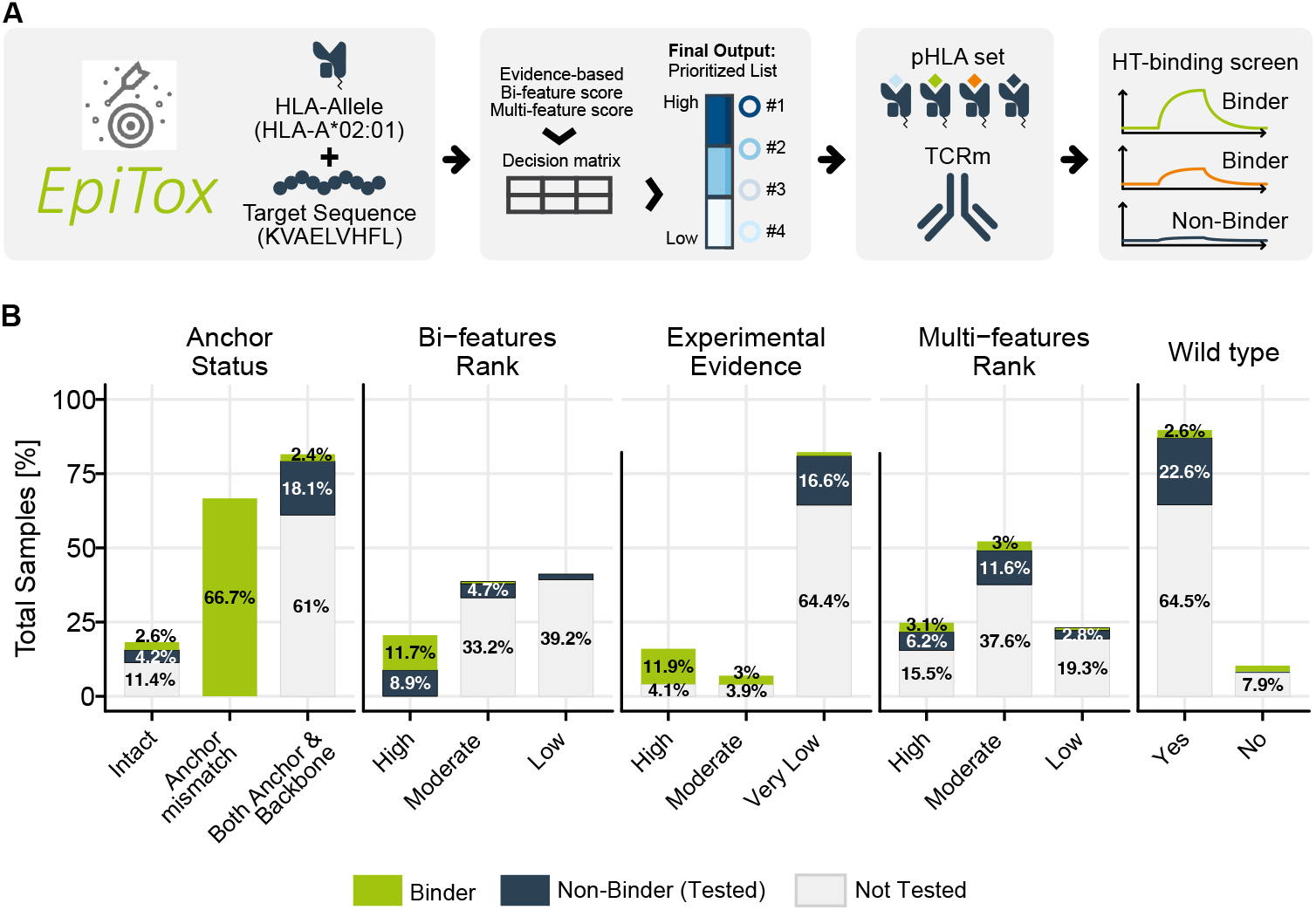
Experimental evaluation of EpiTox predicted peptides: (A) Illustration of off-target prediction and validation. (B) Proportions of predicted versus tested peptides across different classification categories.

#### MAGEA3 as a test case for EpiTox-based TCRm off-target risk assessment

To demonstrate EpiTox utility for comprehensive off-target safety evaluation, we tested a publicly available MAGEA3-KVAELVHFL CAR (C1511 (27)) in the Fab format for binding against a selection of EpiTox-predicted off-targets using our proprietary high-throughput interaction analysis (highSCORE) (Fig. 5.A, Methods 5). While this validation assessed antibody-specific cross-reactivity rather than universal tool performance, results demonstrate that EpiTox identifies physiologically plausible off-targets.

In total, 538 peptides were tested using our highSCORE assay, including the target, exceeding typical experimental validation scales in computational immunology studies. Among these, 171 peptides containing exactly one mismatch served dual purposes: their predicted affinity to HLA-A*2:01 values were used as input for the TPT method (Methods 2), while their experimentally measured binding affinities were used for antibody binding profile characterization. Six of these 171 peptides had already been identified as off-target candidates in FindTopes’ initial list of 1454 putative off-targets. Fig. 5.B illustrates the experimental validation outcomes from our 373-peptide test set. The test set captured substantial diversity across our prediction framework. The pool included Bi-feature classifications (n = 299, 80% High risk), reflecting the peptide selection strategy at the time of experimental design. Despite this numerical bias, the set included sufficient representation from Multi-feature classifications (n = 135, 36% High risk) and Contextual evidence classifications (n = 259, 69% very low risk) to evaluate the complementary information provided by each scoring approach. Testing 373 predicted peptides identified 36 crossreactive with antibody C1511 (9.7% hit rate). Of these, 10 were from the known 18 off-targets, 14 were previously reported in MHC studies and 12 represent novel discoveries. Of 10/18 known off-targets, 8 (80%) were ranked high by both Bi-feature and Multi-feature methods (Table 2). Fig. 6 separates the 36 confirmed binders into known off-targets, left panel, and novel binders, right panel, displaying gene expression profiles of their source gene relative to the target. Known off-targets (n = 10) predominantly showed target-like expression patterns (6/10), while novel off-targets (n = 26) were enriched for elevated expression relative to the target (21/26, ≥2-fold higher).

**Fig. 6.**
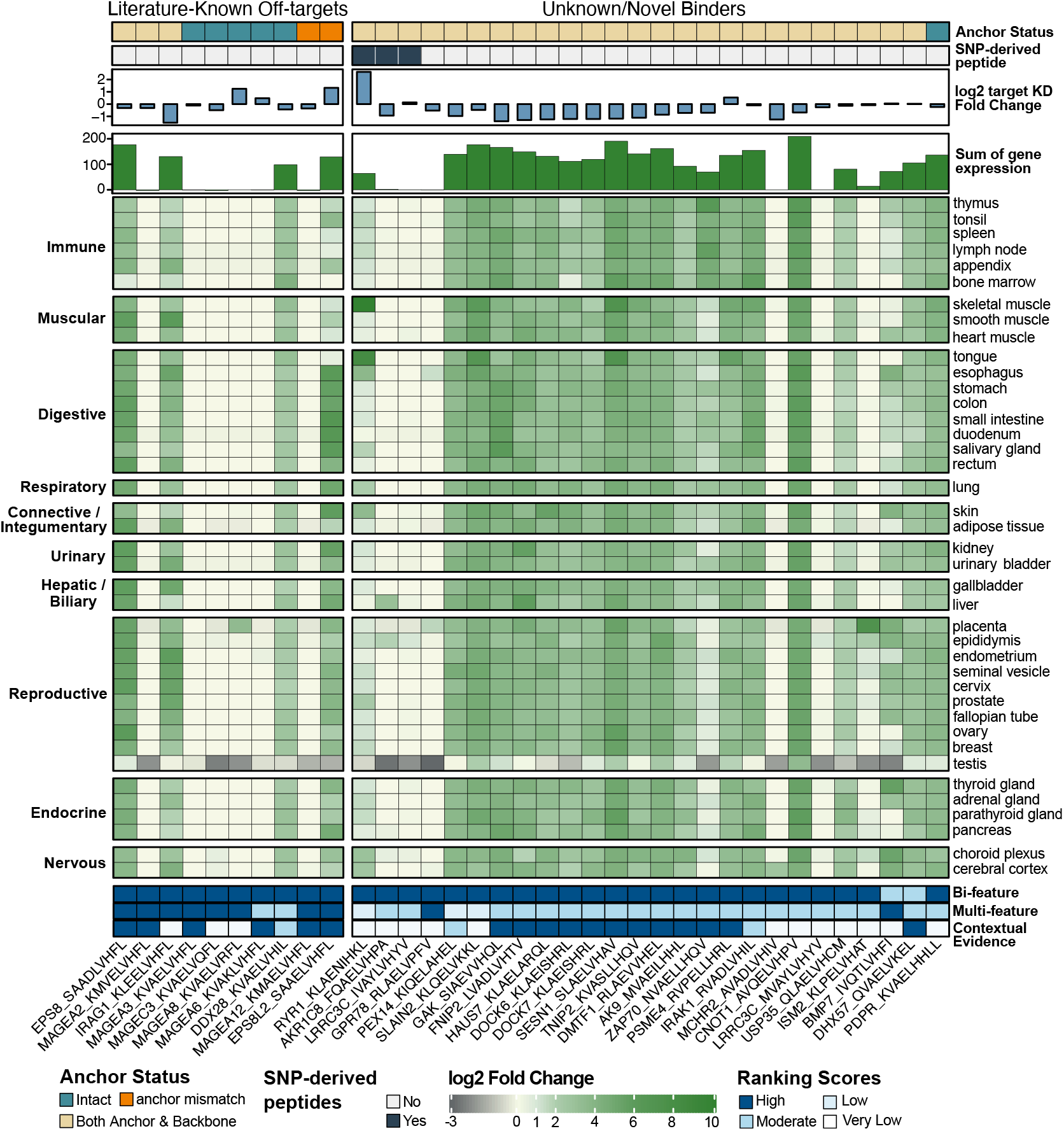
Confirmed binders of EpiTox predicted peptides: Heatmap showing gene expression profiles across multiple tissues for relevant 36 peptides showing crossreactivity. Gene expression values are displayed as log2 fold-change relative to target expression in the tissue. Top annotations from outermost to heatmap: known peptides in MAGEA3 studies & patent versus other; Off-targets mismatch profile at anchor position compared to the target; wildtype versus SNP-derived peptides; log2-transformed KD fold-change relative to target; negative values indicate lower binding affinity compared to target and vice versa; bar plot showing total gene expression summed across tissues per peptide. Bottom annotations: Bi-feature rank, Multi-feature rank, and Experimental Evidence.

#### Population-specific variants reveal novel high-affinity off– targets

We tested 35 SNP-derived peptides representing different score disagreement patterns (all novel, experimental/contextual evidence = very low) (Supplementary Table 5). Three peptides bound the antibody: **Q5T2L2-FQAELVHPA** (rs376200113, p.Arg94Gln) showed high predicted HLA-A*02:01 affinity (KD = 23.4 nM) and BLOSUM62 similarity of 15, compared to its wildtype version (KD = 1584.9 nM, BLOSUM62 = 14). The peptide was classified as high-risk by Bi-feature score and moderate by Multi-feature score. Experimental testing confirmed binding with modest reduction relative to target (log2 KD FC = –0.9). Source protein expression was similar to target, with 4-fold elevation in liver and 2-fold in epididymis. **P21817-KLAENIHKL** (rs746904839, p.Glu879Lys) showed similar predicted affinity to its wildtype version (13 nM) and identical BLOSUM62 score (24). Both versions were classified as high-risk by Bi-feature score and low risk by Multi-feature score. The SNP-derived peptide bound at higher affinity than target (log2 KD FC = 2.6). Source protein showed elevated expression across all 40 normal tissues compared to target, with highest levels in skeletal muscle and tongue (>8 log2 FC). **A6NJW4-IVAYLVHYV** (rs140521451, p.Met236Ile) showed slightly reduced predicted affinity compared to wildtype (22.5 nM vs 16.7 nM) and BLOSUM62 score (23 vs 25). The peptide was classified as high risk by Bi-feature score and moderate by Multi-feature score. Experimental testing showed binding comparable to or elevated compared to the target (log2 KD FC = 0.14). Source protein expression was target-like overall, with elevation in epididymis (log2 FC = 0.9 almost 2-fold higher), choroid plexus (log2 FC = 0.4), and liver, heart muscle and thyroid gland with equal elevated levels (log2 FC = 0.26).

We compared experimentally confirmed binders against the contextual evidence framework established in PrediTopes. Twelve binders were neither found in previously disclosed off-target lists nor in major MHC databases (IEDB, PEPREP), resulting in very low experimental evidence classification (Fig. 6). Three of these were SNP-derived peptides described in the previous section. Among the remaining nine peptides, approximately half showed elevated expression across multiple tissues, while the other half exhibited expression levels comparable to the therapeutic target with tissue-specific elevation in placenta, esophagus, and choroid plexus.

The three scoring approaches showed distinct patterns across the known and unknown peptides. While Bi-feature and Multi-feature methods ranked most cross-reactive peptides as high risk, Contextual evidence scoring predominantly assigned low risk to unknown peptides due to absence of prior empirical data. This divergence was particularly evident in the unknown binders set, where Bi-feature relied on sequence similarity and affinity, Multi-feature incorporated physico-chemical properties, and Contextual evidence reflected literature support. These complementary information sources enabled stratification of cross-reactivity risk based on different biological principles.

## Discussion

Cross-reactivity of TCRm therapeutics against off-targets arises from multiple partially independent factors: sequence similarity, anchor preservation, HLA presentation, physicochemical compatibility and genetic variation (5). **EpiTox** was designed around this multidimensional reality, integrating broad peptide discovery, contextual evidence, and complementary scoring strategies (Fig. 1, Table. 3). Rather than treating complexity as a hurdle, our platform leverages it to reveal patterns of cross-reactivity that simpler approaches would miss. **FindTopes** identified 18 of 19 previously reported peptides/off-targets, validating our approach (Fig. 2.B, Supplementary Table 3). Target position template (TPT) mismatch analysis showed that 19% of off-targets retained intact anchor positions (Fig. 2.C), consistent with predicted HLA complex formation. Cases such as MAGEA12 and EPS8L2 underscore the biological relevance of anchor-preserved sequences, while PPP2R1B-GIAELVHFS illustrates how mismatches exclusively at anchor positions compromise binding (KD = 1313.3 nM, was not bound by the antibody).

In practice, this multidimensional approach was evident in our experimental screening of a anti-MAGEA3 TCRm against 538 different peptide HLA complexes (Fig. 5.B). 171 selected for antibody binding profile characterization and 373 representing our heterogeneous off-target discovery pool, with some overlap between the two sets. From the 373 peptides in our discovery pool, 36 (9.7%) were confirmed cross-reactive to this specific antibody, including 10 previously reported off-targets (Fig. 6, left panel). The multidimensional scoring framework captures cross-reactivity complexity through complementary metrics rather than reducing it to a single score (Table 3). Bi-feature score captured most binders, reflecting high sequence similarity and predicted HLA affinity. Multi-feature score identified a subset of peptides with strong physicochemical similarity, while Evidence-based was absent or low for many novel candidates (Fig. 6 right panel), as expected due to lack of prior experimental data. It is important to emphasize that off-target binding is TCRm specific and that binding profiles change based on the tested TCRm.

Notably, only 5 of 36 binders ranked high across all three metrics, whereas many binders exhibited divergence among dimensions. This divergence illustrates that true off-targets can manifest through distinct mechanisms, sequence conservation, physiochemical mimicry, genetic variation or evidence prior evidence information, proving that multidimensional assessment is necessary to capture the full landscape. For instance, 12/36 novel peptides with low contextual evidence were still validated experimentally as potential physiologically relevant off-targets, demonstrating the predictive value of the three metrics, even when prior data are unavailable.

Genetic variants are rarely integrated systematically into offtarget discovery, leaving a critical gap in identifying risks before clinical manifestation (Table 3). We screened 150 SNP-derived peptides (10% of the pool), the first empirical assessment of SNPs in TCRm off-target screening (Table 3). Most of the SNP-derived peptides (83%) contained backbone or anchor alterations, while 17% retained intact anchors similar to the overall predicted pool (Fig. 2.C & Fig. 6). Experimental evaluation of 35 SNP-derived peptides identified three binders, two with antibody affinities exceeding the target peptide (see Results: 5). These off-targets would only manifest in individuals carrying specific SNPs, highlighting how early SNP integration informs antibody optimization and enables safer, population-conscious therapeutics.

Our evidence framework further addresses data incompleteness and MHC database bias by assigning posterior probabilities rather than binary inclusion/exclusion (Table 3). This approach differentiates identical sequences in distinct protein contexts (e.g., MAGEA3 vs. MAGEA9, 5 Supplementary Table 7), revealing biologically meaningful differences that sequence-based (and structural) methods alone would miss. Experimental validation confirmed the value of this probabilistic approach: 15 of 36 binders (41.7%) originated from the very low evidence tier (23.08% posterior probability), including three previously reported MAGEA3 off-targets (Fig. 6, left panel) and 12 putative novel discoveries, three of which were SNP-derived peptides (Fig. 6, right panel). This demonstrates convergence between two independent bias-mitigation strategies: systematic SNP integration and Bayesian handling of data incompleteness.

These results highlight four key features: [1] calibrated uncertainty without dismissal, [2] discovery potential from under-studied peptides, [3] added value beyond database-only methods, and [4] a modular architecture that is easily extendable to incorporate additional data sources or types (Fig. 3).

Overall, the framework appropriately balances high confidence in experimentally supported peptides (High tier: 99.74%) with cautious but non-dismissive evaluation of low-evidence peptides, enabling both precision and discovery. Across modules, the multidimensional scoring strategy revealed a consistent pattern: peptides confirmed experimentally often scored high in at least one risk dimension, while divergence among metrics highlighted context-dependent vulnerabilities. For example, Q15546-KVVELFFYL showed strong Bi-feature and experimental evidence but low Multifeature, whereas Q9UII4-LVAELVGYR exhibited the opposite profile (Fig. 6). These cases illustrate that no single metric captures the full spectrum of cross-reactivity, and that systematic combination uncovers subtle patterns of risk. These results demonstrate EpiTox’s ability to capture crossreactivity, although they do not guarantee universal performance. Validation using an independent TCRm targeting a different antigen MAGEA4 (28) confirmed consistent identification of experimentally verified off-targets by EpiTox, supporting the broader applicability of the approach. However, even with flexible search parameters, sequence-based off-target identification suffers from a founder effect: peptides that are highly dissimilar and fall beyond predefined similarity thresholds will be missed. In the same study, EpiPredict, a TCRm-specific machine learning framework trained on high-throughput kinetic off-target screening data, addresses this limitation by predicting functionally relevant, sequence-unrelated off-targets. Both EpiTox and EpiPredict are integrated into ValidaTe, a comprehensive and unbiased framework for assessing off-target risk in TCRm therapeutics (manuscript in preparation).

## Conclusions

TCRm cross-reactivity arises from multiple partially independent factors including sequence similarity, anchor preservation, HLA presentation, physicochemical compatibility, and genetic variation. **EpiTox** addresses this multidimensional challenge through integrated scoring strategies that reveal cross-reactivity patterns missed by single-metric approaches.

Experimental validation demonstrated the framework’s practical value. Screening 538 peptide-HLA complexes against an anti-MAGEA3 TCRm identified 36 binders, including 26 novel candidates. Only 5 of 36 binders ranked high across all three metrics, confirming that off-targets manifest through distinct mechanisms and that multidimensional assessment is essential for comprehensive risk capture.

Three key advances distinguish EpiTox: systematic SNP integration, contextual evidence handling and multidimensional scoring. Screening 150 SNP-derived peptides identified three binders, two exceeding target affinity, revealing population-specific risks. Meanwhile, 40.5% of confirmed binders originated from the very low evidence tier, demonstrating that calibrated uncertainty enables discovery from understudied peptides without sacrificing precision.

While sequence-based searches introduce inherent limitations, validation with an independent TCRm confirmed consistent off-target identification. EpiTox is integrated into ValidaTe, which addresses the founder effect through complementary prediction of sequence-unrelated off-targets. By combining systematic peptide discovery, genetic variation, and multidimensional scoring, EpiTox enables more complete risk assessment and supports development of safer TCRm therapeutics.

## ACKNOWLEDGEMENTS

AI use: GPT-4, GPT-5 and Claude Sonnet 4 models were employed exclusively for language refinement during manuscript preparation.

## Author contributions statement

H.A.*, S.S., and O.S. developed the conceptual design and overall architecture of the EpiTox workflow. The implementation, development, and integration of the computational pipeline were carried out by H.A., O.S., and S.S. The design and construction of the supporting PEPREP database were performed by M.A. and H.A. The MAGEA3 study was designed by S.S., K.H., H.A., and J.B. Experimental validation and verification of computational results, including highSCORE-based measurements, were conducted by T.H., P.M., P.H., K.H., A.S., and G.R. H.A., S.K., A.S., O.S., J.B., and S.S. wrote the manuscript. All authors contributed to manuscript review and revisions.

## Supplementary data

As provided.

## Footnotes

1 https://gnomad.broadinstitute.org/news/2017-02-the-genome-aggregation-database/

