## supplementary data, tables and figures for "EpiTox: A Multi-Modular Framework for Population-Aware Off-Target Prediction Highlighting MAGEA3 Cross-Reactivity"

### Materials

#### 1. Data sources and acquisition

Table 1: dataset, access date and other info

| Dataset | Purpose | Access date | Filtering | Source |
|---|---|---|---|---|
| Uniprot | Human Protein sequences + protein annotation | Feb 2025 | Reviewed sequences + proteins with valid Uniprot ids | https://www.uniprot.org/ |
| gnomAD v2.1.1 leftover to GRCh 38 | SNPs | 2025 | SNPs in exon regions; coding SNPs; AF >0.1; Filter = PASS; population = EUR* | https://gnomad.broadinstitute.org/ |
| Ensembl v110 | VEP indexed cache | 2025 | https://ftp.ensembl.org/pub/release-110/variation/indexed_vep_cache/homo_sapiens_vep_110_GRCh38.tar.gz | |
| HPA v24 | HPA RNA dataset | 2025 | Valid cross-reference with Uniprot-ensembl IDs | https://www.proteinatlas.org/download/rna_tissue_hpa.tsv.zip |
| GTEx v8 | Transcript-level expression | 2025 | | https://storage.googleapis.com/adult-gtex/bulk-gex/v8/rna-seq/GTEx_Analysis_2017-06-05_v8_RSEMv1.3.0_transcript_tpm.gct.gz |
| IEDB | Peptides annotations | 2025 | Human; linear peptide; peptide sequence | https://www.iedb.org/ |
| | term = … (list of peptide sequences)<br>base_uri='https://query-api.iedb.org'<br>search_params = {'linear_sequence': term,<br>'structure_type': 'eq.Linear peptide',<br>'host_organism_iri': 'eq.NCBITaxon:9606',<br>'source_organism_iri': 'eq.NCBITaxon:9606',<br>'order': 'linear_sequence', 'offset': 0} | | | |
| PEPREP | Peptides annotations | 2025 | Human; peptide sequence | BioCopy GmbH |

Table 2: tools and packages that were used directly in EpiTox

| Tools/package | Purpose | Reference |
|---|---|---|
| BLASTp | Sequence search | Mahram et al., *ACM (TRETS)* , 2015 |
| PEPMatch | Sequence search | Marrama et al., *BMC bioinformatics*, 2023 |
| tabix 1.9 | SNPs extraction* | Danecek et al., *Bioinformatics*, 2021 |
| Bcftools | | |
| vep | SNPs annotation | Hunt et al., *Human mutation*, 2022 |
| NetMHCpan | HLA binding predictions | Reynisson et al., *Nucleic Acids Research*, 2020 |
| MHCflurry | | O'Donnell et al., *Cell Systems*, 2020 |
| R packages | | |
| ggplot2 | Visualization | H. Wickham. ggplot2: Elegant Graphics for Data Analysis. Springer-Verlag New York, 2016. |
| lessR | | Gerbing D (2025). lessR: Less Code, More Results. R package version 4.4.5, <https://CRAN.R-project.org/package=lessR>.<br><br>Gerbing DW (2021). "Enhancement of the Command-Line Environment for use in the Introductory Statistics Course and Beyond."<br><br>Journal of Statistics and Data Science Education, 29(3), 251-256. doi:10.1080/26939169.2021.1999871<br><br><https://doi.org/10.1080/26939169.2021.1999871>. |
| biomaRt | Cross-reference and gene annotation | Mapping identifiers for the integration of genomic datasets with the R/Bioconductor package biomaRt. Steffen Durinck, Paul<br><br>T. Spellman, Ewan Birney and Wolfgang Huber, Nature Protocols 4, 1184-1191 (2009).<br><br>BioMart and Bioconductor: a powerful link between biological databases and microarray data analysis. Steffen Durinck, Yves<br><br>Moreau, Arek Kasprzyk, Sean Davis, Bart De Moor, Alvis Brazma and Wolfgang Huber, Bioinformatics 21, 3439-3440 (2005). |
| rmarkdown | Report generation | Allaire J, Xie Y, Dervieux C, McPherson J, Luraschi J, Ushey K, Atkins A, Wickham H, Cheng J, Chang W, Iannone R (2024).<br><br>rmarkdown: Dynamic Documents for R. R package version 2.29, <https://github.com/rstudio/rmarkdown>. |
| tidyr | *Data manipulation and analysis* | Wickham H, Vaughan D, Girlich M (2024). tidyr: Tidy Messy Data. R package version 1.3.1,<br><br><https://CRAN.R-project.org/package=tidyr>. |
| Python modules | | |

| | | |
|---|---|---|
| Panda | *Data manipulation and analysis* | *Wes McKinney,* Proceedings of the Python in Science Conference*, 2010* |
| scikit-learn (sklearn) | *Machine learning algorithms and utilities* | *Pedregosa et al.,* Journal of Machine Learning Research*, 2011* |
| NumPy | *Numerical computing, arrays, linear algebra* | *Harris et al.,* Nature*, 2020* |
| Bio.SeqUtils (Biopython) | *Sequence utility functions (molecular weight, composition, etc.)* | *Cock et al.,* Bioinformatics*, 2009* |
| SciPy | *Scientific computing, optimization, statistics* | *Virtanen et al.,* Nature Methods*, 2020* |
| peptides | *Peptide analysis (hydrophobicity, pI, composition, etc.)* | *Wirth & Nystrom,* Journal of Open Source Software*, 2020* |
| Framework | | |
| R 4.4.1 | R Core Team (2024). _R: A Language and Environment for Statistical Computing_. R Foundation for Statistical Computing, Vienna, Austria. <https://www.R-project.org/>. | |
| Python 3.9.16 | Van Rossum, G., & Drake, F. L. (2009). Python 3 Reference Manual. Scotts Valley, CA: CreateSpace. | |
| Conda | | |
| snakemake 7.24.2 | Mölder, F., Jablonski, K.P., Letcher, B., Hall, M.B., Tomkins-Tinch, C.H., Sochat, V., Forster, J., Lee, S., Twardziok, S.O., Kanitz, A., Wilm, A., Holtgrewe, M., Rahmann, S., Nahnsen, S., Köster, J., 2021. Sustainable data analysis with Snakemake. F1000Res 10, 33. | |
| **we followed gnomAD population's description for selection (see https://gnomad.broadinstitute.org/news/2017-02-the-genome-aggregation-database/)* | | |

#### 2. PEPREP: in-house database for pHLA annotations

Along with public resources, EpiTox uses in-house established resource for experimentally validated pHLA complexes, peptides repertoire (PEPREP). PEPREP aggregates and harmonizes public and propriety data to provide different information for each pHLA, for example HLA-allele, disease, and tissue.

#### 3. Known peptides/off-targets in MAGE-A3 context

Three studies and one patent were used to create a list of 18 peptides that was used as disclosed or known off-targets for MAGE-A3 case. The peptides sequences and annotations were incorporated as they were provided by the papers.

Table 3: known peptides in MAGE-A3 context as speculated or confirmed off-targets

| Gene name | peptide | mismatch | source |
|---|---|---|---|
| MAGEA9, MAGEA3 | KVAELVHFL | 0 | PMID23377668, PMID21149604, WO2021173674A1 |

| | | | |
|---|---|---|---|
| DDX28 | KVAELVHIL | 1 | PMID23377668, WO2021173674A1, PMID27439771 |
| MAGEA12 | KMAELVHFL | 1 | PMID23377668, PMID21149604, WO2021173674A1, PMID27439771 |
| MAGEA6 | KVAKLVHFL | 1 | PMID21149604, WO2021173674A1 |
| MAGEA8 | KVAELVRFL | 1 | PMID21149604, WO2021173674A1 |
| MAGEC2 | KVAELVEFL | 1 | PMID21149604 |
| MAGEC3 | KVAELVQFL | 1 | PMID23377668, WO2021173674A1, PMID27439771 |
| EPS8L2 | SAAELVHFL | 2 | WO2021173674A1, PMID27439771 |
| MAGEA1 | KVADLVGFL | 2 | PMID21149604, WO2021173674A1 |
| MAGEA2 | KMVELVHFL | 2 | PMID21149604, WO2021173674A1 |
| MAGEA4 | KVDELAHFL | 2 | PMID21149604, WO2021173674A1 |
| MAGEA5 | KVADLIHFL | 2 | WO2021173674A1 |
| MAGEB18 | KVVSLVHFL | 2 | PMID23377668, WO2021173674A1 |
| MRV11, MRVI1 | KLEELVHFL | 2 | PMID23377668, WO2021173674A1, PMID27439771 |
| EPS8 | SAADLVHFL | 3 | PMID23377668, WO2021173674A1, PMID27439771 |
| MAGEA10 | KVTDLVQFL | 3 | WO2021173674A1 |
| MAGEF1 | TVAELVQFL | 3 | PMID23377668, WO2021173674A1 |
| PPP2R1B | GIAELVHFS | 3 | PMID23377668, WO2021173674A1 |
| MAGEA11 | KIIDLVHLL | 4 | WO2021173674A1 |

### 4. EpiTox architecture

#### 4.1 PrediTopes: HLA binding prediction and experimental evidence

##### 4.1.1 Data structure

While quantitative binding measurements exist for some peptides (e.g., IC50 binding affinities, raw MS values), categorical evidence (validation: yes/no; binding: strong/intermediate/weak; class: I/II) provides more complete coverage across the dataset. We therefore utilized categorical evidence as the primary input, producing discrete posterior probabilities when evidence combinations recur. The framework accommodates future integration of quantitative data where available.

Furthermore, experimental data were aggregated for each protein-peptide pair (uniprot id – peptide sequence), preserving allele-binding relationships and associated metadata including tissue/disease context, binding affinity measurements, and study quality indicators. This protein-specific aggregation ensures that evidence assessments reflect the exact biological context of each peptide-source.

Table 4: Likelihood ratios used in Bayesian framework

| | |
|---|---|
| **Evidence quality:** | |
| Multiple high-relevance studies: target allele + tissue context, | LR = 12 |
| Single high-relevance study | LR = 8 |
| Multiple/single medium-relevance studies | LR = 6/4 |
| Multiple/single low-relevance studies | LR = 3/2 |
| No experimental evidence | LR = 0.3 (conservative penalty) |
| **HLA allele specificity:** | |
| Target allele match | LR = 10 |
| Different allele | LR = 2 |
| HLA class match (when allele is unknown): | Same: LR = 2<br>Different: LR =0.5 |
| **Binding affinity:** | |
| Base LR: strong = 5, intermediate = 3, weak = 2, non-binder = 1 | |
| Target-specific multipliers: strong = 2.5x, intermediate = 1.5x, weak = 1x, negative = 0.2x | |
| **Tissue/disease context:** | |
| Normal tissue | LR = 10 |
| Same disease (optional) | LR = 6 |
| Similar tissue | LR = 3 |

#### Supplementary figure 1

LR sensitivity analysis

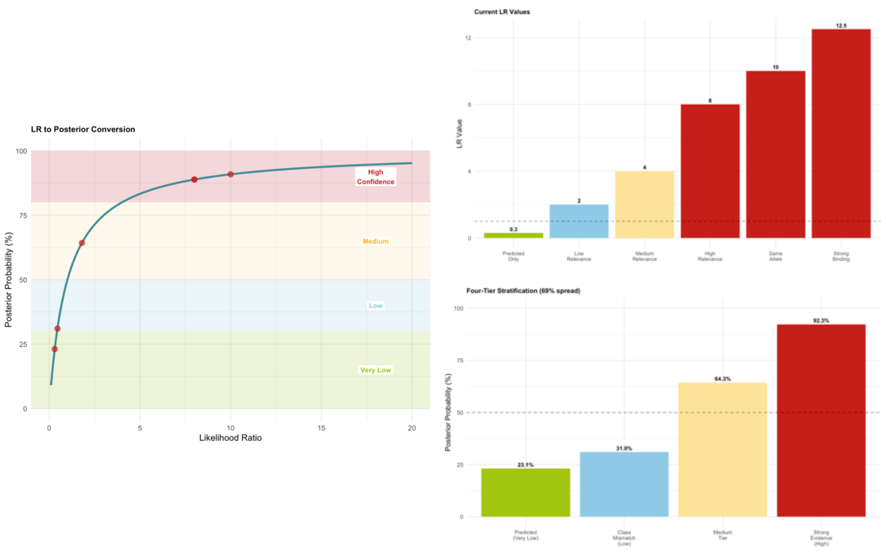

### Results and methods

Table 5: Position-specific HLA-binding of MAGEA3

| Position | AA | Class | FC_mean | Sensitivity | % Better than target |
|---|---|---|---|---|---|
| P1 | K | Classical | 5.96 | 39 | 15.79 |
| P2 | V | Classical | 24.41 | 108.67 | 15.79 |
| P3 | A | Permissive | 1.63 | 3.9 | 31.58 |
| P4 | E | Neutral | 1.91 | 2.8 | 5.26 |
| P5 | L | Permissive | 1.12 | 0.6 | 31.58 |
| P6 | V | Neutral | 1.43 | 1.46 | 15.79 |
| P7 | H | Neutral | 1.62 | 2.65 | 0 |
| P8 | F | Neutral | 1.5 | 1.8 | 5.27 |
| P9 | L | Classical | 305.99 | 1135.26 | 10.53 |

*AA: Target sequence amino acid wise.*
*FC_mean: mean fold change in KD upon substitution.*
*Sensitivity: normalized range of KD change.*
*% Better than WT: percentage of substitutions with better KD than target.*

#### Supplementary figure 2

Target positional template (TPT)

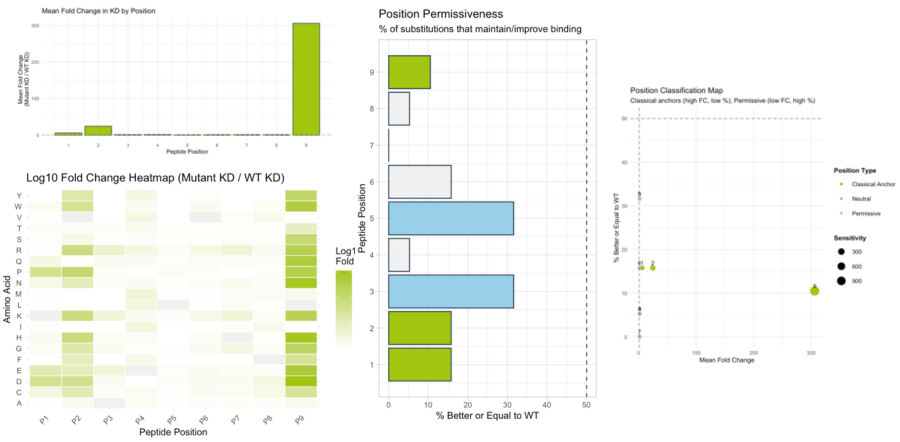

Table 6: Identified off-target candidates from different MAGE families (A, B, C, F & E)

| uniprot | Gene Name | Peptide | Blosum62 similarity | Edit distance | Wildtype | HLA-A*2:01 Affinity |
|---|---|---|---|---|---|---|
| | | | **MAGEA family** | | | |
| **P43357** | **MAGEA3** | **KVAELVHFL** | **44** | **0** | **Yes** | **14,3961329** |
| P43362 | MAGEA9 | KVAELVHFL | 44 | 0 | Yes | 14,3961329 |
| P43365 | MAGEA12 | KMAELVHFL | 41 | 1 | Yes | 10,8250373 |
| P43361 | MAGEA8 | KVAELVRFL | 36 | 1 | Yes | 52,6518232 |
| P43360 | MAGEA6 | KVAKLVHFL | 40 | 1 | Yes | 23,7568242 |
| P43356 | MAGEA2 | KMVELVHFL | 37 | 2 | Yes | 10,885883 |
| P43355 | MAGEA1 | KVADLVGFL | 31 | 2 | Yes | 36,2984481 |
| P43358 | MAGEA4 | KVDELAHFL | 34 | 2 | Yes | 29,9630256 |
| P43363 | MAGEA10 | KVTDLVQFL | 29 | 3 | Yes | 30,3629588 |
| P43364 | MAGEA11 | KIIDLVHLL | 29 | 4 | Yes | 15,3669415 |
| Q7L5Y9 | MAEA (PIG5) | VVAELEKTL | 14 | 4 | Yes | 1046,11964 |
| Q7L5Y9 | MAEA (PIG5) | VVAELQKTL | 14 | 4 | No | 967,514188 |
| | | | **MAGEB family** | | | |
| Q96M61 | MAGEB18 | KVVSLVHFL | 35 | 2 | Yes | 19,2521481 |
| A8MXT2 | MAGEB17 | KTGELVQFL | 28 | 3 | Yes | 104,838719 |
| A2A368 | MAGEB16 | KVAFLVNFM | 27 | 3 | Yes | 1158,76239 |
| Q96LZ2 | MAGEB10 | KVIILVHYL | 28 | 3 | Yes | 38,8062522 |
| O15479 | MAGEB2 | KSGSLVQFL | 21 | 4 | Yes | 1673,43194 |
| O15481 | MAGEB4 | KTKMLVQFL | 20 | 4 | Yes | 7170,88466 |
| O15480 | MAGEB3 | KTNMLVQFL | 19 | 4 | Yes | 1665,05912 |
| Q9BZ81 | MAGEB5 | KVGLLEQFL | 18 | 4 | Yes | 2413,8697 |
| A8MXT2 | MAGEB17 | KVLEFVAKL | 16 | 4 | Yes | 20,0630444 |
| | | | **MAGEC family** | | | |
| Q9UBF1 | MAGEC2 | KVAELVEFL | 36 | 1 | Yes | 19,868702 |
| Q8TD91 | MAGEC3 | KVAELVQFL | 36 | 1 | Yes | 20,4910337 |
| O60732 | MAGEC1 | KVDELARFL | 26 | 3 | Yes | 111,874657 |
| Q8TD91 | MAGEC3 | KVDKLVQFL | 26 | 3 | Yes | 69,3400635 |
| | | | **MAGEF family** | | | |
| Q9HAY2 | MAGEF1 | TVAELVQFL | 30 | 2 | Yes | 40,0224415 |
| | | | **MAGEE family** | | | |
| Q9HCI5 | MAGEE1 | NVAELLQFL | 28 | 3 | Yes | 57,7469841 |

**Kommentiert [HAH1]:** is this needed in the main text or could be shifted to supplementary?

**Kommentiert [SS1R2]:** supplementary

#### Supplementary figure 3

Position-specific mismatches vs. predicted binding affinity

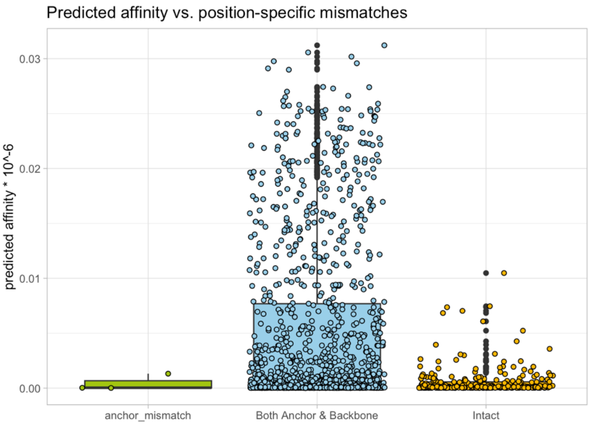


#### Supplementary figure 4

Multi-features ranking score

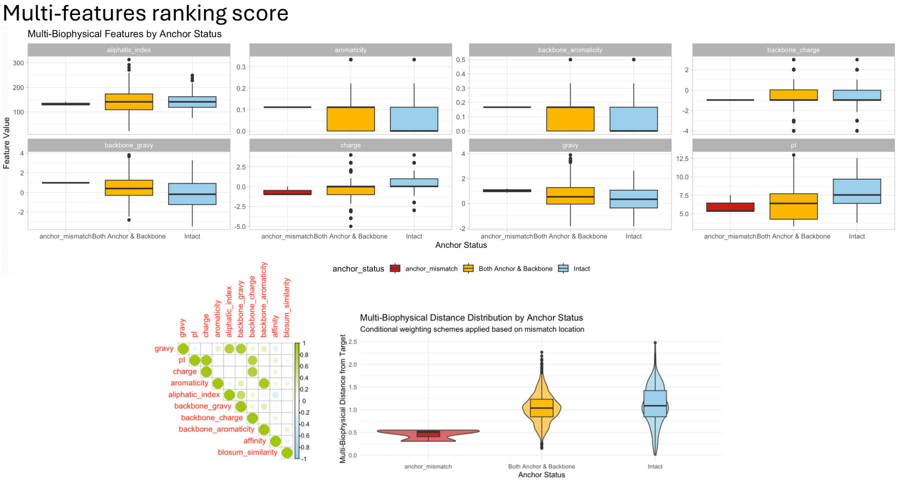

### Supplementary figure 5

Bi-features vs. multi-features assessment

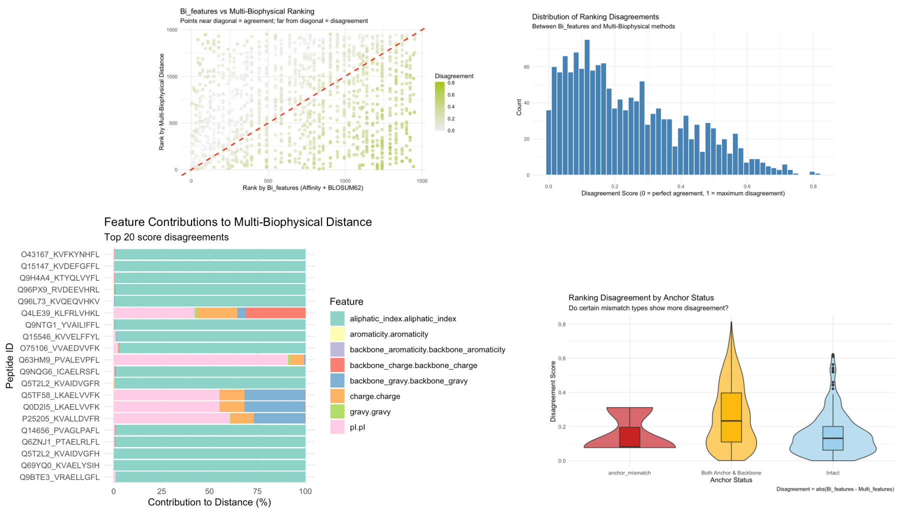


*Table 5: SNP-derived peptide ranked by different scores*

| **Multi-features** | **Bi-features** | | |
|---|---|---|---|
| 2-4 | High | Moderate | Low |
| High | 9 | 15 | 2 |
| Moderate | 1 | 4 | 1 |
| Low | 0 | 3 | 0 |
